# Dissociable Effects of Early and Adolescent Adversity on Emotional Contagion

**DOI:** 10.64898/2026.01.29.702530

**Authors:** Paloma P Maldonado, Erica Berretta, Viviana Canicatti, Xiaoyi Feng, Efe Soyman, Lenca Cuturela, Rajeev Rajendran, Maryam Sadeghi, Akos Babiczky, Georg Goebel, Harm J Krugers, Christian Keysers, Valeria Gazzola

**Author notes:** Contributed equally to the role of first author (Paloma P Maldonado, Erica Berretta). Contributed equally to the role of senior author (Harm J Krugers, Christian Keysers, Valeria Gazzola).

## Abstract

**Background:** Early-life adversity can alter emotional and social development and increase vulnerability to later life stress. We investigated how early adverse experiences (EAE) and later adverse experiences (LAE) shape adult emotional contagion (EC) responses in female and male rats.

**Methods:** EAE was induced using the limited bedding and nesting model during the first postnatal week. LAE was induced via footshocks during adolescence. In adulthood, male and female rats underwent an EC test in which observers witnessed a conspecific receiving footshocks.

**Results:** Adolescence-footshock exposed observers showed cingulate cortex-associated increased immobility, proximity, and attention toward distressed conspecifics during adulthood, compared to adult-exposed and sham animals, both in male and female animals. While EAE did alter maternal care, pup stress physiology, and pup weight, we found evidence that it did not alter immobility during EC. However, female demonstrators paired with EAE observers showed increased immobility, linked to a reduced rate and lower frequency of the observers’ 50-kHz vocalizations. Mediation analysis revealed that a shift toward lower-frequency 50-kHz vocalizations specifically mediated this effect, suggesting a sex-specific pathway by which early adversity shapes social behavior.

**Conclusions:** Early and adolescent adversity influenced distinct aspects of emotional contagion: EAE mediated an observer-to-demonstrator emotional transfer during EC, while LAE impacted a demonstrator-to-observer transfer, with no evidence of additive effects. Our results highlight developmentally specific and sex-dependent mechanisms by which early and later adversity alter social-affective responses in adulthood.

## INTRODUCTION

Traumatic experiences, defined as events involving actual or perceived threats to life, physical safety, or sexual integrity^1^ can significantly shape how humans respond to others’ distress. These responses range from heightened empathy, which may lead to emotional overwhelm, to reduced empathy, which may serve as a protective mechanism^2–4^. The mechanisms underlying this variability are not fully understood, though early life experiences are increasingly recognized as key modulators.

Adverse experiences during early development, collectively termed early-life adversity (ELA), include not only trauma affecting physical integrity but also parental loss, neglect, poverty, and exposure to violence. These experiences activate multiple systems, including the stress response system, particularly the hypothalamic-pituitary-adrenal (HPA) axis, and have been associated with long-term emotional and social outcomes, including the modulation of empathic responses to others^5,6^. Nevertheless, a recent meta-analysis^7^ reported mixed findings regarding the nature of this modulation: 33% of studies linked ELA to increased empathy, 44% to decreased empathy, and 27% found no effect, suggesting that additional factors shape empathic responses.

One critical factor affecting empathy may be mother-infant interactions, often disrupted by ELA. In rodents and humans, the HPA axis is less reactive during early life, a “stress hyporesponsive period”; however, maternal care disruption is one of the few factors that can impair this protective mechanism^8,9^, underscoring its importance. John Bowlby’s “44 Thieves” study^10^ highlighted the link between ELA and empathy, showing that most thief children with “affectionless psychopathy” had experienced prolonged maternal separation. Modern studies confirm this association: children raised in institutional settings exhibit higher levels of callous-unemotional traits^11^.

ELA rarely occurs in isolation; instead, it often increases the risk of additional adversity later in life, a process known as stress proliferation. Individuals exposed to childhood abuse or neglect are significantly more likely to encounter negative life events later in life compared to those without such histories^12–14^. This concept has been linked to a higher likelihood of developing psychopathology, such as depression^12,15^. Supporting this, the two-hit hypothesis in animals^16^ and the stress sensitization model in humans^17,18^ suggest that early stress creates a latent vulnerability that may only manifest behaviorally when triggered by later adversity. This helps explain why ELA increases the risk for mental disorders^19–21^, even though such experiences are neither necessary nor sufficient causes on their own^16,22^. This framework is particularly relevant to empathy, as reduced empathic capacity is frequently observed in individuals with psychopathology^23–25^.

This study investigated how two forms of adversity, early adverse experience (EAE) and a later adverse experience (LAE), affect emotional contagion responses in rats (Figure 1). EAE was modelled using the limited bedding and nesting (LBN) paradigm, which is considered to mimic low socio-economic status, disrupts mother-pup interactions, and activates the HPA axis. LAE was modelled through footshocks delivered either during adolescence or adulthood, allowing the assessment of developmental timing effects on emotional contagion (EC). EC was assessed using a paradigm in which female or male observer rats witnessed same-sex conspecifics receiving footshocks. By using Bayesian statistics^26^, we examined whether we could find evidence for or against main effects of EAE, main effects of LAE, or an interaction between EAE and LAE on behavioral indices of emotional contagion. To examine whether the effects may depend on sex, we further examined the effect of sex in same-sex dyads. Finally, as cingulate area 24 has been shown to be necessary for EC in rats and mice^27–29^, and contains mirror neurons activated during both pain experience and pain observation in rats^27^, we asked whether individual differences in c-Fos activation in area 24 are associated with individual difference in EC.

**Figure 1.**
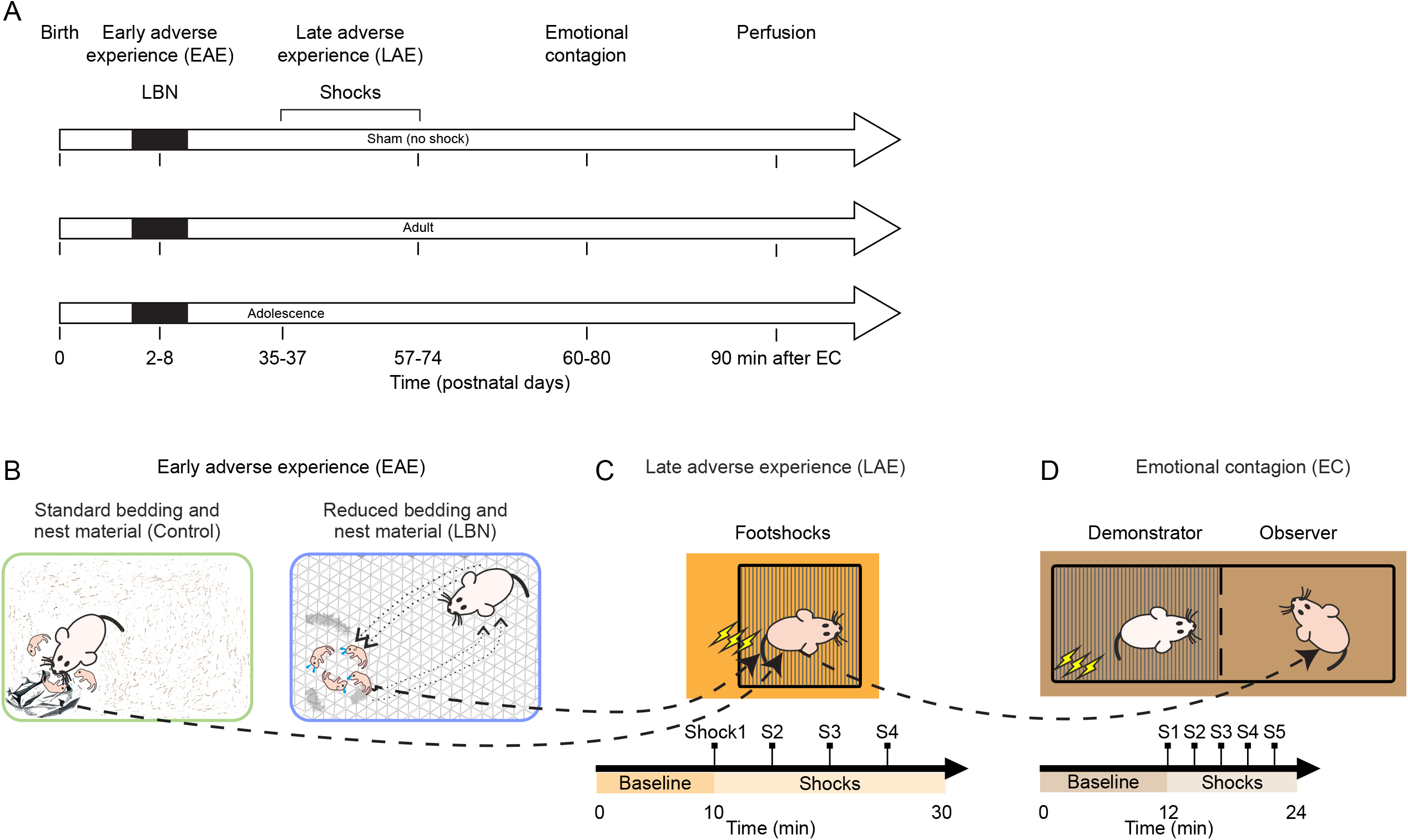
Experimental procedures. A. Time course of the experimental procedures. B. Schematic depiction of the early adverse experience (EAE) protocol. C. Top: schematic depiction of the late adverse experience (LAE) protocol. Bottom: time course of the LAE procedure. D. Top: schematic depiction of the emotional contagion (EC) paradigm. Bottom: time course of the EC procedure. Note, observers are the pups raised in the LBN condition.

## METHODS AND MATERIALS

Extended methods and materials are described in the Supplementary Material.

### Subjects

All experimental procedures were approved by the institutional animal care and use committee of the Royal Netherlands Academy of Arts and Sciences and were in agreement with the European Community Directive 2010/63/EU.

### Experimental procedures

#### LBN

LBN protocol was implemented as described by Molet et al.^30^.

#### LAE procedure

At either adolescent (P35–P37) or adult (P57–P79) age, observer animals underwent a footshock or a sham procedure.

#### Emotional contagion test

The habituation and handling procedures related to the EC test are described in Supplementary method figure 1. EC test was performed as described previously^27,28,31^.

### Analysis

#### Maternal behaviors

One-hour video recordings of maternal behaviors were manually scored using the open-source software BORIS (v.7.7.3, Italy^32^).

#### Entropy

Entropy refers to a measure of ‘randomness’ or the ‘unpredictability’ of an entire random variable. Shannon entropy was used to compute the entropy index measuring the unpredictability of a single dam’s behavioural transitions from P2-P8^33–35^.

#### Immobility

Immobility was manually scored using the open-source software BORIS. Periods of no visibility during the EC test were excluded when estimating immobility durations.

#### Vocalizations

Vocalizations from both the LAE and EC test were analyzed using DeepSqueak (version 2.6.1, via MATLAB R2019a).

#### Pose estimation

DeepLabCut was used to track the observer and demonstrator rats separately after the video was cropped in half.

#### Cell counting

Cell quantification was done using raw multichannel fluorescent images. We used a custom made whole-brain cell quantifier (AMBIA^36^). Sections were matched to a standard rat brain atlas^37^ to ensure anatomical consistency across samples.

### Statistical analysis

All data are shown as mean ± SEM, except for the frequency of 50-kHz vocalizations, which was reported as mode ± SEM. Statistical analyses were performed using JASP^38^.

## RESULTS

### Developmental timing of LAE shapes EC behavior

Rats were subjected to an emotional contagion (EC) test between P60 and P80, in which they observed an unfamiliar, age- and sex-matched conspecific (the demonstrator), receiving footshocks (Figure 1A and D). EC refers to the transfer or sharing of affective states between individuals, often occurring through observation of another’s distress, a phenomen well-described in both rats and mice^39,40^. Figure 2A, depicts the temporal profile of observer’s immobility. In our experimental design, footshocks were applied as a within-subjects factor, while EAE, LAE and sex were treated as between-subjects factors. The between-subjects factors included the following levels: for EAE, control and LBN conditions; for LAE, sham, adult, adolescence (ado) groups; and for sex, female and male. The within-subject factor shocksOBS had five levels (shock 1 to 5). Consistent with Han et al.^31^, female observers displayed significantly less immobility than males (repeated-measures ANOVA, EAE X LAE X sex X shocksOBS, main effect of sex: F(1, 95) = 13.819, p = 3.4e-4, BFincl = 57.36). Notably, both female and male rats exposed to footshocks during adolescence showed significantly increased immobility during the EC compared to those exposed in adulthood or those in the sham group (repeated-measures ANOVA, EAE X LAE X sex X shocksOBS, main effect of LAE: F(2, 95) = 22.117, p = 1.3e-8, BFincl > 1000; post hoc Holm test ado vs adult p = 1.4e-5, ado vs sham p = 1.1e-8). Importantly, we found evidence against a main effect of EAE on observer immobility (F(1, 95) = 0.094, p = 0.76, BFincl = 0.048), and evidence against an interaction effect of EAE X LAE (F(2,95) = 0.479, p = 0.621, BFincl = 0.055), suggesting that EAE did not alter EC as proxied by observer immobility, nor did it alter the response to later adversity. To explore the neural correlates of these behavioral effects, we examined c-Fos expression in area 24 following the EC test, which has previously been shown to be necessary for vicarious freezing in rats^27,28^. We observed that the c-Fos density was positively correlated with individual differences in immobility, but only in the group exposed to footshocks during adolescence (Figure 2B, right).

**Figure 2.**
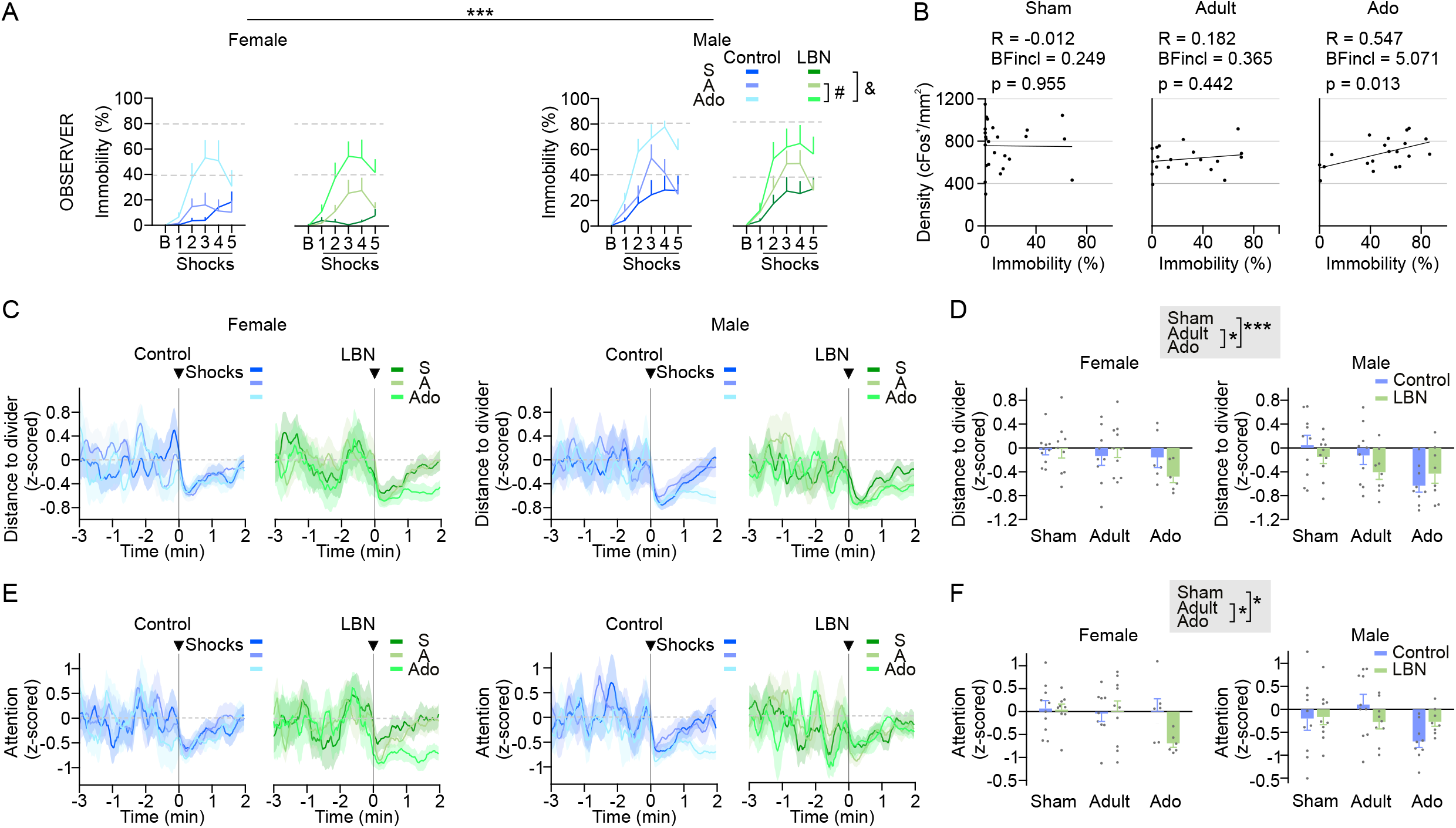
Female and male observers immobilized more during EC, directed their attention towards the demonstrators, and approached the divider between the observers and the demonstrators more when the LAE occurred during adolescence. A. Percentage of immobility for baseline (B) and five-shock periods in the control condition and the LBN condition for female (left) and male (right) observers. Female control and LBN Sham n = 10, adult n = 10, adolescent n = 8. Male control Sham n = 10, adult n = 10, adolescent n = 10; LBN Sham n = 10, adult n = 11, adolescent n = 8. ***p = 3.4e-4. #p = 1.4e-5. &p = 1.1e-8. B. Correlation between c-Fos expression and percentage of observer immobility after EC. Because no significant differences were observed between control and LBN conditions, or between sexes, all animals were pooled for this analysis. Sham n = 25, adult n = 20, adolescence n = 20. C. Time course of the z-scored distance between the head of the observer and the divider for control and LBN females (left) and males (right). The shock period corresponds to the average of the first two-minutes of the five shocks. Female control Sham n = 10, adult n = 9, adolescent n = 6; LBN Sham n = 9, adult n = 10, adolescent n = 6. Male control Sham n = 10, adult n = 10, adolescent n = 9; LBN Sham n = 10, adult n = 9, adolescent n = 8. D. Average z-scored distances between the observer’s head and the divider during the last 10 seconds of the shock periods for females and males. Adolescence vs adult *p = 0.03; ado vs sham ***p = 7.75e-4. E. Time course of the z-scored attention (degrees) between the head of the observer and the head of the demonstrator for control and LBN females (left) and males (right). The shock period corresponds to the average of the first two-minutes of the five shocks. F. Average z-scored angle between the observer and demonstrator’s head during the last 10 seconds of the shock periods for females and males. Adolescence vs adult *p = 0.041; ado vs sham *p = 0.037.

Witnessing shocks to a demonstrator does not only trigger immobility, but also approach and orienting to the demonstrator^27^. Female and male rats exposed to LAE during their adolescence approached the divider between the observer and the demonstrator significantly more compared to those animals exposed during their adulthood and animals in the sham group, reflected in negative z-score values (Figure 2C and D, ANOVA, EAE X LAE X sex, main effect of LAE, F(2,94) = 7.233, p = 0.001, BFincl = 20.441, post hoc Tukey test ado vs adult p = 0.041, ado vs sham p = 7.75e-4). Similar to immobility, approach also showed evidence against an effect of EAE (F(1,94) = 0.94, p = 0.335, BFincl = 0.125). Right after each shock, observers also turned their heads towards the demonstrators, with both female and male observers that experienced LAE during their adolescence turning significantly more compared to those that experienced it during their adulthood and those in the sham group (Figure 2E and F, ANOVA, EAE X LAE X sex, main effect of LAE: F(2,94) = 4.122, p = 0.019, BFincl = 1.273, post hoc Tukey test ado vs adult p = 0.03, ado vs sham p = 0.037). Again, we found evidence against an effect of EAE (F(1,94) = 0.788, p = 0.377, BFincl = 0.111) and EAE X LAE (F(2,94) = 0.241, p = 0.786, BFincl = 0.067) on head orientation.

### EAE changes female 50-kHz vocalizations affecting demonstrator behavior

Following this evidence for a lack of EAE effect on immobility, proximity, and orienting during EC, we turned to the behavior of the demonstrators, who were unfamiliar with the observers, as prior work has demonstrated that the state of the observer can influence the response of the demonstrator in shock observation paradigms^28^. As expected, demonstrators exhibited increased immobility after footshocks (Figure 3A). Across the shock periods, male demonstrators displayed more immobility than females (Figure 3B, repeated-measures ANOVA, EAE X LAE X sex X shocksDEM, main effect of sex: F(1, 95) = 11.613, p = 9.6e-4, BFincl = 3.733), with evidence against a main effect of EAE condition (F(1, 95) = 0.804, p = 0.372, BFincl = 0.04), LAE (F(2, 95) = 0.063, p = 0.939, BFincl = 0.007) or interaction effect of EAE X LAE (F(2, 95) = 1.393, p = 0.253, BFincl = 0.003). However, when analyzing the total immobility across the full shock session, we observed a sex-specific effect of EAE. Female demonstrators paired with EAE observers showed significantly higher total immobility compared to those paired to control female observers, while this effect was absent in males (Figure 3C, ANOVA, EAE X LAE X sex, interaction effect EAE X sex: F(1,103) = 7.346, p = 0.008, BFincl = 8.370, post hoc Holm test, female EAE vs female control: p = 0.004, BFincl = 9.975; male EAE vs male control p = 0.987, BFincl = 0.295).

**Figure 3.**
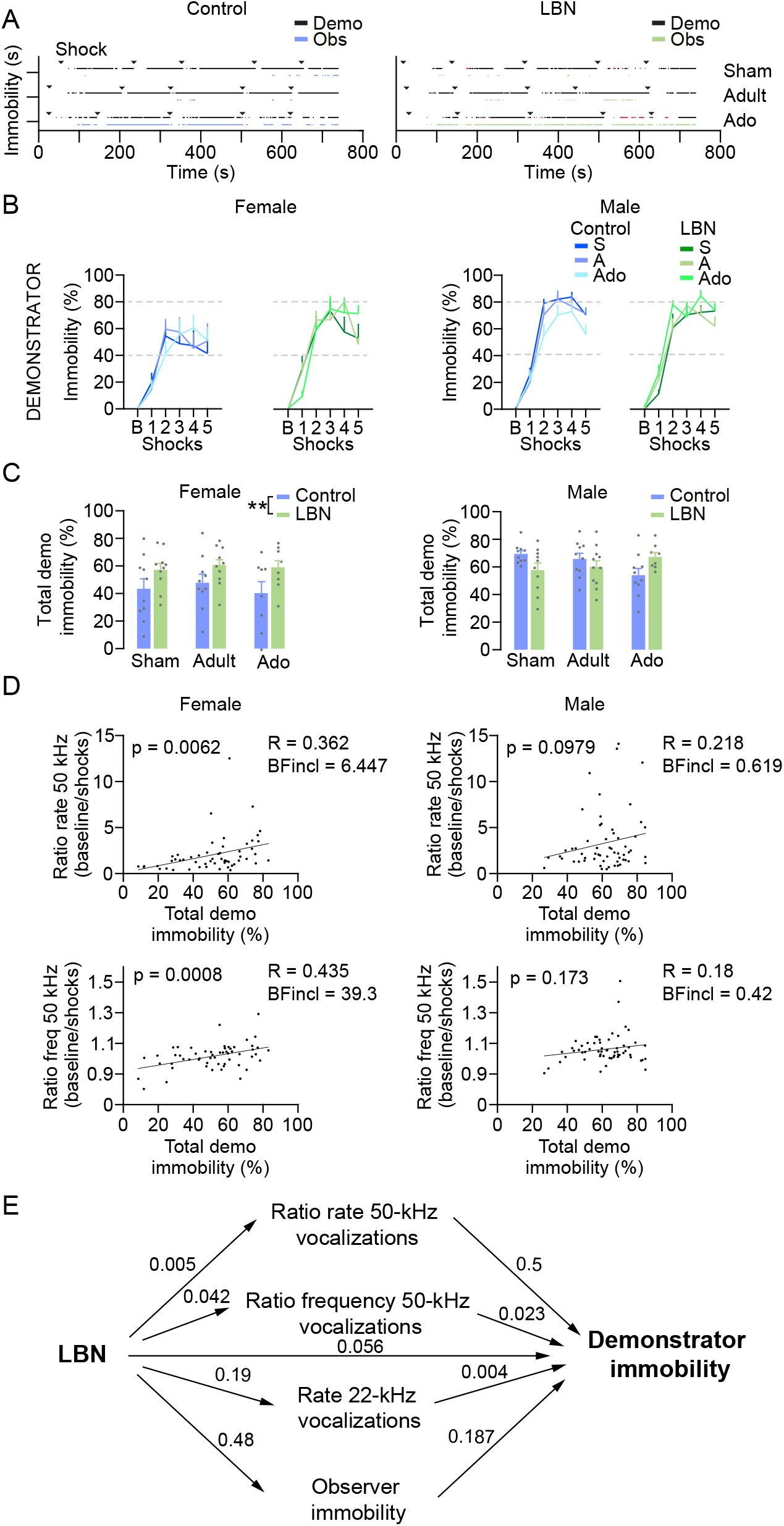
Female demonstrators immobilize more when paired with LBN than with control observers. A. Time course of immobility of 3 females from sham, adult, and adolescent LAE in the control (left) and the LBN condition (right). Red colour indicates periods in which the demonstrator’s immobility was not possible to quantify. The triangle symbol indicates the timing of the shocks. B. Percentage of immobility for baseline (B) and 5 shock periods in the control condition and the LBN condition for female (left) and male demonstrators (right). Female control and LBN Sham n = 10, adult n = 10, adolescent n = 8. Male control sham n = 10, adult n = 10, adolescent n = 10; LBN Sham n = 10, adult n = 11, adolescent n = 8. C. Percentage of total demonstrator immobility during the 5 shock periods for females (left) and males (right) in control and LBN conditions. **p = 0.004. D. Top: the correlation between the ratio of 50-kHz vocalization rate and total immobility of demonstrators from control and LBN conditions and the three LAE groups for females (left) and males (right). Bottom, correlation between the ratio of 50-kHz vocalization frequency and total immobility of demonstrators from control and LBN conditions and the three LAE groups for females (left) and males (right). E. Mediation model. Numbers correspond to p-values.

Since demonstrators had neither undergone LBN nor been housed with LBN animals, the observed differences in their immobility—based on the EAE condition of their observers— suggest that female observers raised under LBN conditions may have emitted signals during the EC test that influenced demonstrator behavior, specifically increasing immobility. Furthermore, our results from Figure 2 indicated that female observer immobility, proximity, and orientation was not affected by LBN, ruling out the possibility that increased demonstrator immobility resulted from mirroring an observer’s immobility or due to systematically altered proximity or attention by their observers. We therefore investigated whether vocalizations produced by female observers during the intershock intervals might have served as a communicative cue influencing demonstrator behavior. In our set up, a single microphone was positioned at the center of the emotional contagion apparatus, and vocalizations could therefore not be confidently attributed to the observer or demonstrator. However, we analyzed overall vocalizations to infer group-level effects.

We first examined the 22-kHz vocalizations. 22-kHz vocalizations, typically emitted seconds after footshocks, are associated with negative affective states occurring during adverse situations^41^. No such vocalizations were emitted during the baseline period. 22-kHz vocalizations were emitted during the intershock intervals (ISI), although they were not significantly affected by either EAE condition (Supplementary figure 1A, ANOVA, EAE X LAE: F(1,50) = 1.65, p = 0.205, BFincl = 0.384) or LAE (Supplementary figure 1A, ANOVA, EAE X LAE: F(2,50) = 1.091, p = 0.344, BFincl = 0.233). We then focused on the higher frequency vocalizations conventionally referred to as “50 kHz” vocalizations. 50-kHz vocalizations are emitted by both juvenile and adult rats and are often emitted in appetitive and rewarding contexts^41^. Overall, there was a significant reduction in 50-kHz vocalization rate during the ISI periods compared to baseline (Supplementary figure 1B-D, ANOVA, EAE X LAE X shocks, main effect of shocks: F(1,100) = 6.114, p = 0.015, BFincl = 1.497). Importantly, this reduction was modulated by the EAE condition, as shown by a significant EAE X shocks interaction (Supplementary figure 1D, ANOVA, EAE X LAE X shocks, interaction effect EAE X shocks: F(1,100) = 4.424, p = 0.038, BFincl = 0.611). This was further confirmed by comparing the ratio of rates between baseline and ISI periods (Supplementary figure 1E, ANOVA, LBN X LAE, main effect of EAE: F(1,50) = 6.374, p = 0.015, BFincl = 2.768), with evidence of absence of an effect of LAE (Supplementary figure 1E, ANOVA, EAE X LAE, main effect of LAE: F(2,50) = 0.539, p = 0.587, BFincl = 0.188). Because ultrasonic vocalization frequency can convey emotional valence or arousal levels^42^, we next examined this parameter. The frequency was measured at the point of maximum intensity. We observed a significant downward shift in the frequency of 50-kHz-vocalizations during the ISI compared to baseline (Supplementary figure 1B-C and F, ANOVA, EAE X LAE X shocks, interaction effect EAE X shocks: F(1,100) = 6.386, p = 0.013, BFincl = 1.116). Again, this effect was supported by the baseline/shock frequency ratio (Supplementary figure 1G, ANOVA, EAE X LAE, main effect of EAE: F(1,50) = 10.293, p = 0.002, BFincl = 13.174), with evidence of absence for a LAE effect (Supplementary figure 1G, ANOVA, EAE X LAE, main effect of LAE: F(2,50) = 0.215, p = 0.807, BFincl = 0.236).

Together, these results suggest that changes in both the rate and frequency of 50-kHz vocalizations emitted during the shock observation period by female EAE observers that experienced EAE differed from those that did not and might have contributed to the increased immobility reported in demonstrators paired with EAE observers. To explore this association, we assessed the relationship between observer vocalizations and demonstrator behavior by calculating correlations between the baseline-to-shock ratio of 50-kHz vocalization rate and frequency, and the percentage of immobility displayed by their demonstrators. This analysis revealed a significant correlation in females (Figure 3D, see statistical details in figure legend), indicating that in female dyads, lower 50-kHz vocalization rate and frequency during shock observation were associated with increased demonstrator immobility. No significant correlations were observed in male pairs. To further integrate our findings, we computed a mediation analysis, which showed that within the parameters we measured, the lowering of the frequency of the 50-kHz vocalization during shock observation, captured by the ratio of 50-kHz vocalizations frequencies, measured in EAE female dyads best explains and mediates the increased immobility of EAE-paired demonstrators (Figure 3E, see statistical details in figure legend).

### EAE-induced changes in maternal behavior and pup physiology without corresponding increase in corticosterone levels

One possible explanation for the lack of EAE effects in the EC test is that our LBN condition may not have sufficiently altered maternal behavior. To explore this, we examined behavioral features and outcomes previously reported for this model. The LBN condition was implemented from postnatal day (P) 2 until P9 (Figure 1A). Pups reared under LBN condition exhibit significantly lower weight compared to controls, starting from P26 in females and males. This reduction was large in effect size, persisting until the final assessment at P49 (Figure 4A, see statistic details in figure legend). These results replicate previous findings regarding body weight outcomes following LBN exposure^43,44^, and further demonstrate the long-lasting impact of EAE extending weeks beyond the cessation of the manipulation. Both sexes showed reduced weight in the LBN condition, and when expressed as a percentage of bodyweight, no significant sex difference was observed (Figure 4B, repeated-measures ANOVA, age X sex, main effect of sex: F(1, 55) = 3.47; p = 0.068, BFincl = 2.619), but an age X sex interaction effect (F(10, 550) = 2.457, p = 0.007, BFincl = 5.293).

**Figure 4.**
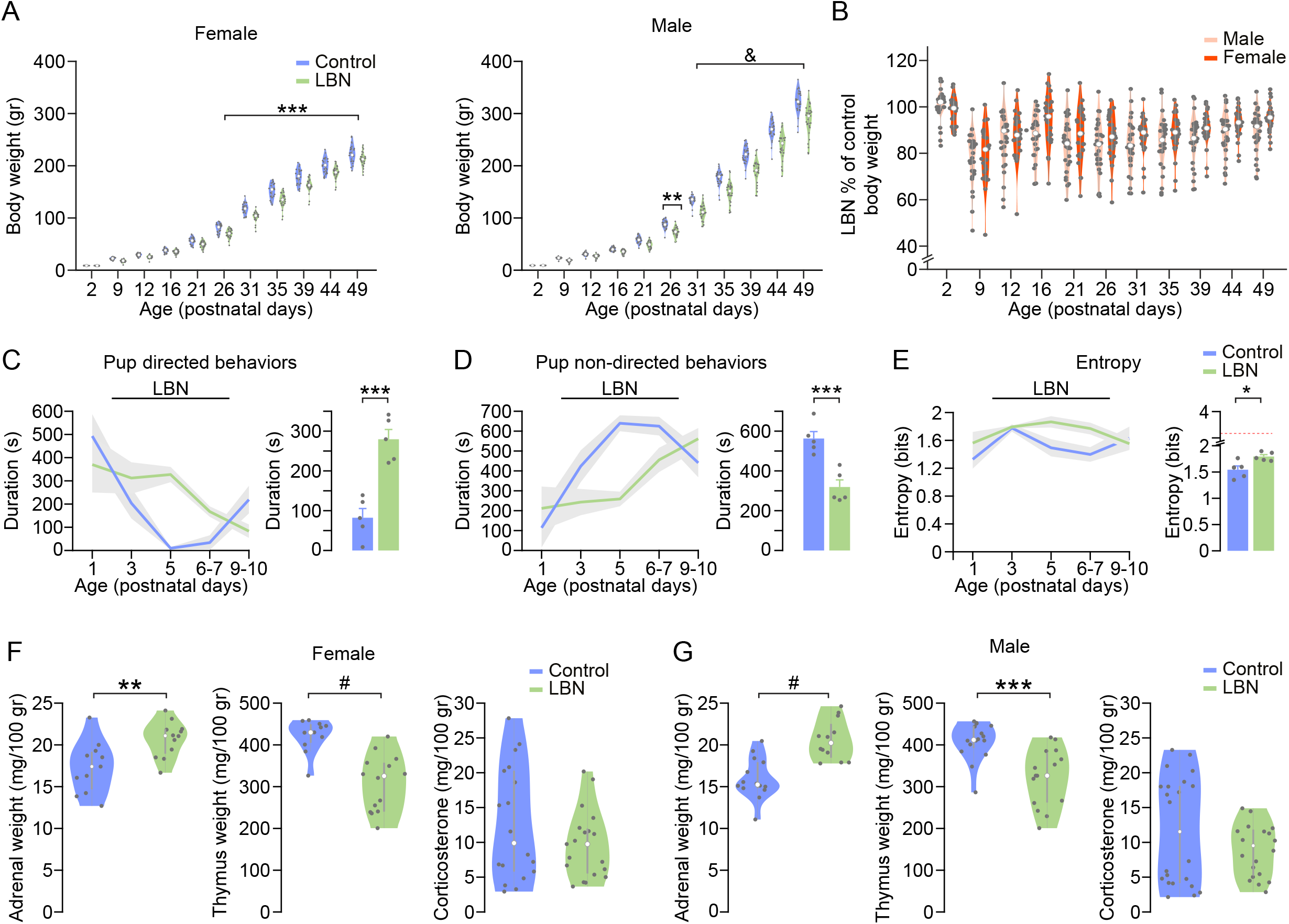
LBN affects maternal behaviors and offspring’s body weight, but not corticosterone levels. A. Female (left) and male (right) weight time courses in control and LBN conditions. Left: control n = 28, LBN n = 28. Repeated-measures ANOVA, LBN x age, main effect of LBN: F(1, 54) = 24.477, p = 7.72e-6, BFincl > 1000. This corresponds to a large effect size (**η**^2^ = 0.312). Interaction effect LBN x age, post hoc Holm test, from P26-P49 all ***p > 0.001. Right: control n = 30, LBN n = 29. Repeated-measures ANOVA, LBN x age, main effect of LBN: F(1, 57) = 42.026, p = 2.303e-8, BFincl > 1000. Interaction effect LBN X age, post hoc Holm test, **p = 0.001, from P31-P49 all &p > 1.4e-7. B. Animal weights in the LBN condition as a percentage of the weights in the control condition. Note that there is no difference between females and males. C. Left: pup-directed maternal behaviour time-course. Right: average duration across all ages in the LBN condition. Every dot is a nest. Control n = 5, LBN n = 5. Mann-Whitney U test. ***p = 0.008, BF_10_ = 2.35. D. Left: non-pup-directed maternal behaviour time course. Right: average duration across all ages in the LBN condition. Every dot is a nest. Control n = 5, LBN n = 5. Mann-Whitney U test. ***p = 0.008, BF_10_ = 2.316. E. Left: entropy time course. Right: average entropy across all ages in the LBN condition. Every dot is a nest. Control n = 5, LBN n = 5. Two-tailed t-test. *p = 0.014, BF_10_ = 4.152. The red dashed line indicates the theoretical maximum entropy (ln2(12)). F. Left: average adrenal weight of female pups raised in control (n = 11) and LBN (n = 12) conditions. **p = 0.006. Middle: average thymus weight of female pups raised in control (n = 11) and LBN (n = 14) conditions. #p = 8.7*10e-5. Right: average corticosterone concentration of female pups raised in control (n = 18) and LBN (n = 19) conditions. G. Left: average adrenal weight of male pups raised in control (n = 14) and LBN (n = 12) conditions. #p = 5.5*10e-5. Middle: average thymus weight of male pups raised in control (n = 16) and LBN (n = 15) conditions. ***p = 3.1*10e-4. Right: average corticosterone concentration of male pups raised in control (n = 22) and LBN (n = 20) conditions.

Pup-directed maternal behaviours (pup licking and grooming, pup carrying, arched-back nursing, low nursing, and passive/side nursing^45^), were significantly altered under LBN condition. LBN dams showed an overall increase in the duration of pup-directed behaviors across the manipulation period (Figure 4C, two-tailed t-test, p = 0.0004, BF_10_ = 55.5). Conversely, non-pup-directed maternal behaviors (self-grooming, off-nest, rearing, eating/drinking), were reduced in LBN dams compared to controls (Figure 4D, two-tailed t-test, p = 0.001, BF_10_ = 21.3). A hallmark of the LBN model is the reduction in the predictability of maternal behaviors^46,47^. In line with previous reports^34,48^, LBN dams exhibited greater unpredictable behaviors, as evidenced by increased entropy compared to controls (Figure 4E, two-tailed t-test, p = 0.011, BF_10_ = 5.0).

As a final step, we examined whether potential stress-related physiological changes, could account for the observed outcomes. Given that the quantification of these markers requires sacrificing the individuals, this was performed in a separate cohort of animals. Pups were sacrificed at P9, immediately following LBN manipulation. In both females and males, LBN-exposed pups displayed significantly higher adrenal weight (Figure 4F, female, two-tailed t-test, p = 0.006, BF_10_ = 8.0; Figure 4G, male, two-tailed t-test, p = 5.5*10e-5, BF_10_ > 300) and decreased thymus weight relative to controls (Figure 4F, female, two-tailed t-test, p = 8.7*10e-5, BF_10_ > 300; Figure 4G, male, two-tailed t-test, p = 3.1*10e-4, BF_10_ = 100.2). However, corticosterone levels did not differ between LBN and control condition (Figure 4F, female, two-tailed t-test, p = 0.2, BF_10_ = 0.625; Figure 4G, male, two-tailed t-test, p = 0.428, BF_10_ = 0.657).

### Pain responses to footshocks are modulated by developmental timing of LAE but not by EAE

Finally, to better understand the difference between experiencing footshocks during adolescence vs adulthood on observer immobility during EC, we analyzed the behavior during the LAE manipulation. Footshocks elicit a range of observable behaviors^40^; here, we focused specifically on immobility and vocalizations.

Immobility during the intershock intervals was significantly higher in animals shocked during adolescence than adulthood (Figure 5A, repeated-measures ANOVA, EAE X LAE X sex X shocks, main effect of LAE: F(1, 59) = 25.088, p = 5.3e-6, BFincl > 1000), and in males compared to females (repeated-measures ANOVA, EAE X LAE X sex X shocks, main effect of sex: F(1, 59) = 20.008, p = 3.6e-5, BFincl = 211.99). The interaction effect between EAE X LAE X sex was weak (repeated-measures ANOVA, EAE X LAE X sex X shocks, F(1, 59) = 4.298, p = 0.043, BFincl = 0.779; post hoc Holm test, male control ado vs male control adult, p = 6.2*10e-4), and nonsignificant if looking at the average immobility across all shocks (ANOVA, EAE X LAE X sex, interaction effect EAE X LAE X sex, F(1, 60) = 3.194, p = 0.079, BFincl = 0.687).

**Figure 5.**
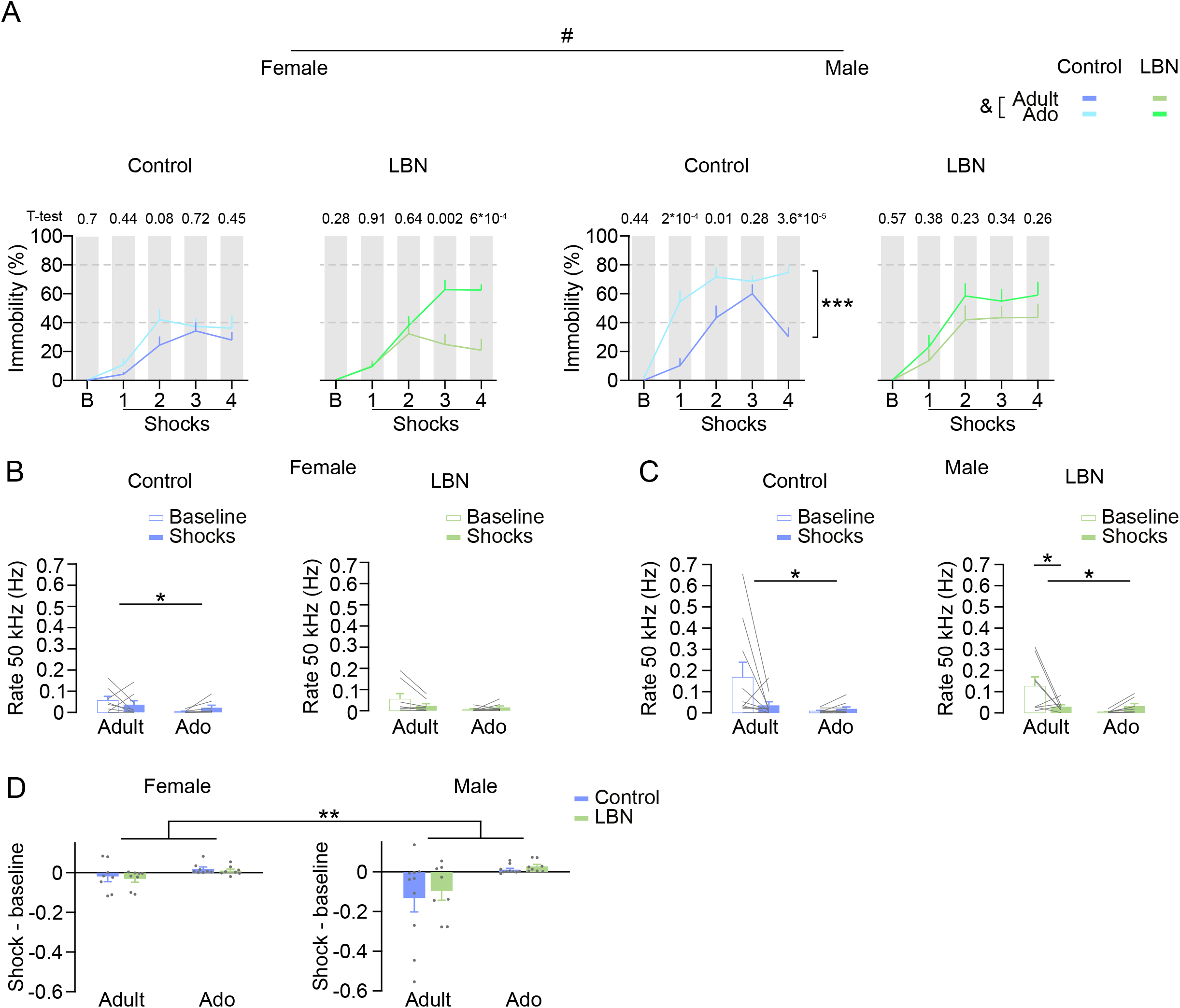
LAE induces more immobility during adolescence than adulthood and changes 50-kHz vocalizations. A. Time course of immobility in control and LBN conditions for females and males during footshock exposure. &p = 5.3e-6. #p = 3.6e-5. ***p = 6.2*10e-4. B. Left: average rate of 50-kHz vocalizations during baseline and four-shock period in the control condition for females. Adult n = 8, adolescent n = 8. ANOVA, shocks X LAE, main effect of LAE: F(1,28) = 4.89, *p = 0.035, BFincl = 1.632. Right: average rate of 50-kHz vocalizations during baseline and four-shock period in the LBN condition for females. Adult n = 8, adolescent n = 8. C. Left: average rate of 50-kHz vocalizations during baseline and four-shock period in the control condition for males. Adult n = 10, adolescent n = 10. ANOVA, shocks X LAE, main effect of LAE: F(1,36) = 5.716, *p = 0.022, BFincl = 2.547. Right: average rate of 50-kHz vocalizations during baseline and four-shock period in the LBN condition for males. Adult n = 8, adolescent n = 8. ANOVA, shocks X LAE, main effect of LAE: F(1,28) = 6.607, *p = 0.016, BFincl = 4.399. Interaction effect shocks X LAE, F(1,28) = 7.170, p = 0.012, BFincl = 4.896, post hoc Tukey test, baseline adult vs shock adult: *p = 0.03. D. Change in 50-kHz vocalization rate (shock – baseline) shown separately for females and males. **p = 0.001.

On average, animals emitted around 20 squeaks, with an average duration of 40 ms, only during the four shock periods, with no significant differences as a function of age of LAE, or EAE (Supplementary figure 2A and B, see statistical details in figure legend). Loudness was not analyzed due to potential variability in microphone positioning.

We then analyzed 22-kHz vocalizations during the ISI and found no significant difference in total duration as a function of LAE (Supplementary figure 2C, ANOVA, EAE X LAE X sex: p = 0.103, BFincl = 0.548) or EAE (Supplementary figure 2C, ANOVA, LBN X LAE X sex: p = 0.932, BFincl = 0.158). However, males emitted significantly longer 22 kHz vocalizations than females (Supplementary figure 2C, ANOVA, EAE X LAE X sex, main effect of sex: F(1,60) = 21.4, p = 2.1e-5, BFincl = 648).

We found a trend indicating that footshock exposure during adulthood reduced 50-kHz vocalization rates relative to baseline (no-shock period), whereas exposure during adolescence led to an *increase* in these vocalizations (Figure 5B and C, see statistic details in figure legend). The quantification of these trends indicated a main effect of LAE (Figure 5D, ANOVA, EAE X LAE X sex, main effect of LAE: F(1,60) = 11.965, p = 0.001, BFincl = 31.477) and evidence against EAE (F(1,60) = 0.096, p = 0.757, BFincl = 0.166) and interaction EAE X LAE (F(1,60) = 0.024, p = 0.877, BFincl = 0.204) for both females and males.

## DISCUSSION

In this study, we aimed to investigate whether early exposure to adverse events can alter emotional contagion in adulthood. Specifically, we examined how rats respond to the distress of conspecifics after experiencing early-life adversity (LBN) and later-life adversity (footshocks during adolescence or early adulthood). Our findings demonstrate that these factors influence distinct aspects of emotional contagion in rats, including sex-specific differences.

### Early-life adversity alone may not be sufficient to impair emotional contagion responses

Our Bayesian analysis provides evidence that our early manipulation, LBN, did not affect emotional contagion responses in a manner indicative of the altered emotional contagion that we had predicted based on prior literature. This evidence of absence of an effect on EC-related immobility was found despite large effect sizes of our LBN manipulation on most of the outcome parameters of LBN previously reported in literature (reviewed in Walker et al.^49^): we replicated reductions in body weight and alterations in the weight of organs involved in immune and endocrine responses to stress^50^. However, we did not observe an increase in corticosterone levels. Literature on the effect of LBN on corticosterone levels is mixed, with some studies reporting increases^44,51,52^ and others reporting no changes at all^53,54^. Nevertheless, many of these studies have identified long-lasting changes in other systems. Supporting this, as pointed out by McLaughlin et al.^55^, some critical outcomes of early-life adversity, such as cortical thinning and impairments in language-related abilities, may occur independently of the HPA axis activation, despite the longstanding association between early trauma and stress axis engagement^56^.

### Negative experiences during adolescence leave a stronger trace than those in adulthood

Recency models suggest that mental health outcomes are most strongly linked to proximal, rather than distal stressor events^57,58^. Supporting this idea, recent traumatic or stressful events have been more strongly associated with psychotic symptoms compared to more distant events^59^. However, in our study, we found an opposite pattern: shock exposure experienced during *adolescence* was associated with higher immobility levels, as well as more approach to and orienting towards the demonstrator during emotional contagion testing in adulthood compared to recent shocks experienced during adulthood (Figure 2)—despite the adolescent shock occurring approximately one month prior, and the adult shock only five days earlier. This finding is compatible with the notion that adolescence may be a sensitive developmental period, a concept supported across multiple domains in the literature^60–62^. Consistent with this, we found differences in immobility during the exposure to shocks itself, with animals exposed to footshocks during adolescence showing more immobility than those exposed to the same footshocks during adulthood (Figure 5A).

This heightened sensitivity may also be a reflection of the enhanced memory encoding capacity observed during adolescence in humans^63^. A wide range of experiences, including autobiographical memories and even mundane events, experienced during adolescence are remembered more vividly later in adulthood. Relevant to our findings, fear extinction learning has been shown to be attenuated in adolescence, both in humans and rodents, likely due to reduced synaptic plasticity in the prefrontal cortex^64^. This may reflect a neurobiological mechanism underlying adolescents’ greater difficulty in recovering from a stressful experience, and by extension, why adversity during adolescence leads to heightened emotional contagion in adulthood despite the reduced recency compared to adult adversity.

### Sex-specific enhanced auditory sensitivity to conspecific distress cues

It has been shown that in EC paradigms, information flows bidirectionally between rats: observers influence demonstrators and demonstrators influence observers^28^. This bidirectional information flow helps explain why, in our study, female demonstrators immobilized more when paired with LBN-reared females, while female LBN observers themselves did not display heightened immobility. When looking for possible mediators of this effect, we identified a shift toward lower-frequency 50-kHz vocalizations in female LBN observers as a mediating factor (Figure 3E), suggesting a role for sound-mediated communication in driving demonstrator responses. Low-frequency vocalizations have been associated with negative affective states^42^, and such frequencies propagate more efficiently through the environment due to reduced absorption, scattering, and greater diffraction compared to high frequencies^65^. These properties make low-frequency vocalizations well-suited for communicating danger to conspecifics^66^. Supporting this, Saito et al.^67^ showed that frequency alone, more than other acoustic features like duration or frequency modulation, provided the most information for discrimination between pleasant and distress calls, supporting how this property perceptually could convey the most for the demonstrators. Interestingly, male LBN observers in our study also exhibited a shift to lower frequencies during shock observation, yet this was not sufficient to induce greater immobility in their male demonstrators. This raises the question of how female LBN observers have a stronger impact on their partners. While keeping in mind the explorative nature of this analysis, and therefore the need to reproduce these results, one may speculate that a possible explanation may lie in the sex-specific sensitivity to vocalizations. Females are specifically endowed with the biological machinery to attend to vocalizations, they retrieve isolated pups to the nest by detecting their ultrasonic vocalizations^68,69^, since as altricial species pups cannot move and signal their distress in another manner. This female capability has been described to be mediated by the oxytocin system^70,71^: oxytocin enhances detection of USVs by disinhibiting the auditory cortex^70,72^. This sex-specific sensitivity is explained by the oxytocin receptor expression being higher in the left compared to the right hemisphere, for both dams and virgin females, and higher compared to males^73^. While these mechanisms were originally described in the context of maternal responses to pup USVs, they likely extend to other social contexts, given oxytocin’s broader role in social behaviors^74,75^.

Considering that we showed that USVs convey rich information about internal affective states, a major limitation of the present study is the inability to differentiate between observer and demonstrator USVs during emotional contagion. This prevented us from resolving the temporal dynamics of vocal communication between the two animals, including when and what determines the shift toward lower-frequency 50-kHz vocalizations. In addition, to determine whether stress-related changes in HPA-axis function may nevertheless have contributed to emotional contagion responses, our study would benefit from assessing corticosterone levels at additional time points and/or quantifying glucocorticoid receptor expression, which may better capture long-term alterations in the stress system.

In conclusion, our findings underscore the importance of both early-life social environments and timing in later stress shaping emotional contagion responses. Importantly, when it comes to emotional contagion, our Bayesian analysis provides moderate evidence that the EAE induced by the LBN paradigm had large effects on body weight and classic measures of LBN effects but did *not* impair nocifensive reactions to the distress of the demonstrator, as indexed by immobility, orienting or proximity, and did *not* alter the effect of later exposure to footshocks, as evidenced by evidence against the existence of an EAE x LAE interaction on the same parameters. In contrast, later stress experienced during adolescence led to lasting alterations in nocifensive indicators of emotional contagion associated with activation in area 24 of the cingulate, consistent with adolescence being a sensitive window for emotional learning. Moreover, subtle shifts in vocalization-mediated communication, particularly in female observers, may mediate the transfer of affective states, potentially shaped by maternal care behaviors.

This publication is part of the project Dutch Brain Interface Initiative (DBI2) with project number 024.005.022 of the research programme Gravitation, which is financed by the Dutch Ministry of Education, Culture and Science (OCW) via the Dutch Research Council (NWO) to VG and CK. Someme (OCENW.XL21.XL21.069) to CK. KNAW 3V Fonds (240-245407) to RR. Guangzhou Elite Scholarship to XF. Brain and Cognition grant to CK, VG and HK.

## According to CRediT

PM: formal analysis, Writing – original draft, Visualization, data curation

EB: Investigation, Conceptualization, Supervision, Project administration, data curation

VC: Investigation

XF: Investigation, formal analysis

ES: Investigation

LC: formal analysis RR: Investigation

MS: software

AB: formal analysis

GG: software

HK: Conceptualization, Supervision, Funding acquisition, Project administration

CK: Conceptualization, Writing - Review & Editing, Supervision, Funding acquisition, Project administration

VG: Conceptualization, Writing - Review & Editing, Supervision, Funding acquisition, Project administration

We thank Tallie Baram and Jessica Bolton for their contributions to the implementation of the LBN protocol. We also acknowledge the students who assisted during the project: Ruben de Klerk, Mariana Edwards, Aikaterini Sfyaki, Steven Voges, and Leonie Schulze. Special thanks to Lorenzo De Angelis and Jeniffer Sanguino Gomez for their support with data analysis implementation, and to Mateo Velez-Fort for his feedback on the manuscript.

The authors reported no biomedical financial interests or potential conflicts of interest.

---

**Supplementary Figure 1.**
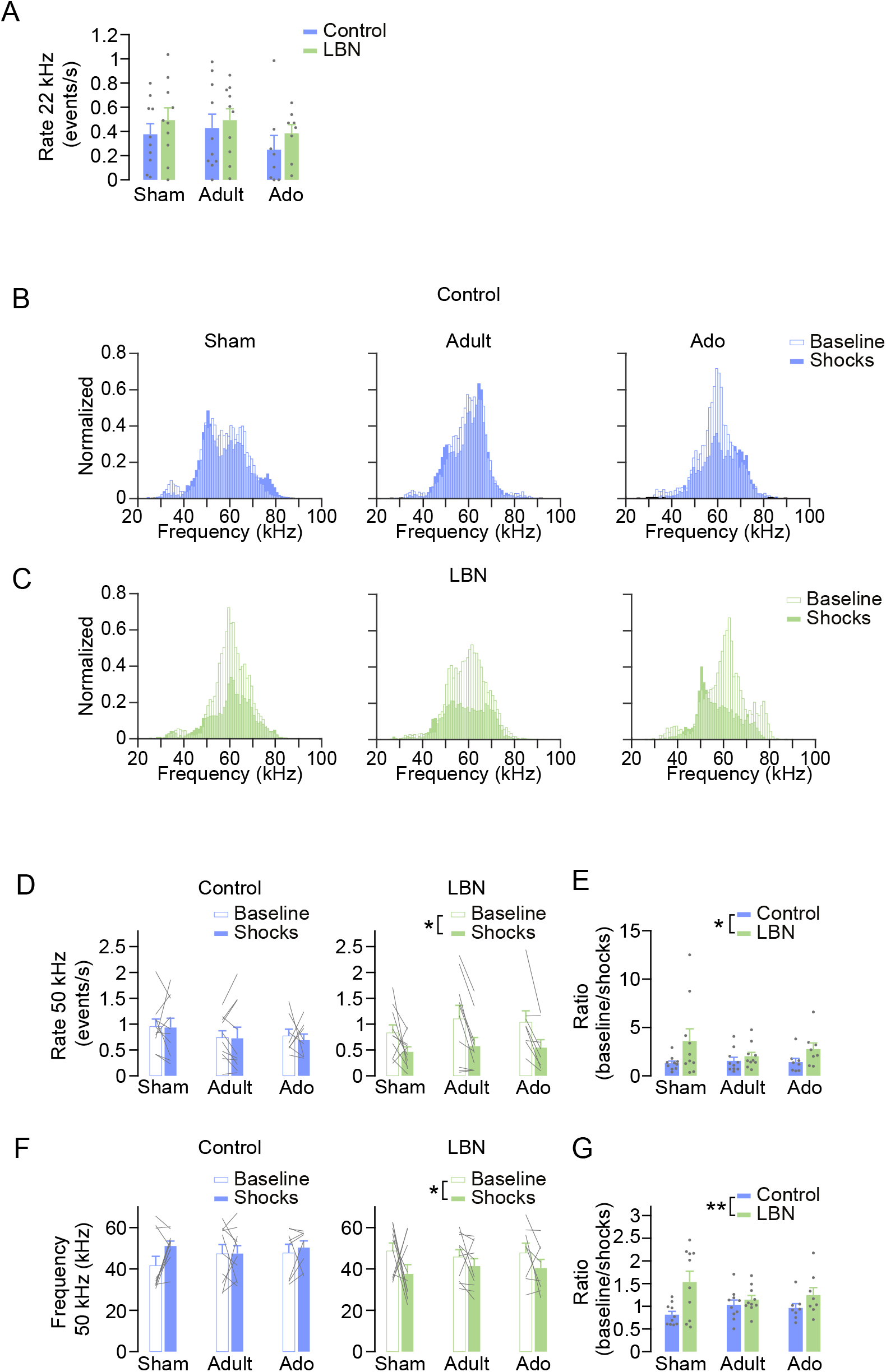
Female 50-kHz vocalizations during EC were affected by EAE, not LAE. A. Average rate of 22-kHz vocalizations during baseline and shock periods in control and LBN females. Control and LBN sham n = 10, adult n = 10, adolescent n = 8. B. Histograms of 50-kHz vocalization frequencies during EC baseline and shock periods in the control condition for sham, adult, and adolescence LAE groups. C. Histograms of 50-kHz vocalization frequencies during EC baseline and shock periods in the LBN condition for sham, adult, and adolescence LAE groups. D. Average rate of 50-kHz female vocalizations during EC baseline and shock periods in the control (left) and the LBN (right) condition. *p = 0.038. E. 50-kHz vocalizations ratio between the average rate during EC baseline and shock periods in control and LBN conditions. *p = 0.015. F. Average frequency of 50-kHz female vocalizations during EC baseline and shock periods in the control (left) and the LBN (right) condition. *p = 0.013. G. 50-kHz vocalizations ratio between the average frequency during EC baseline and shock periods in control and LBN conditions. **p = 0.002.

**Supplementary Figure 2.**
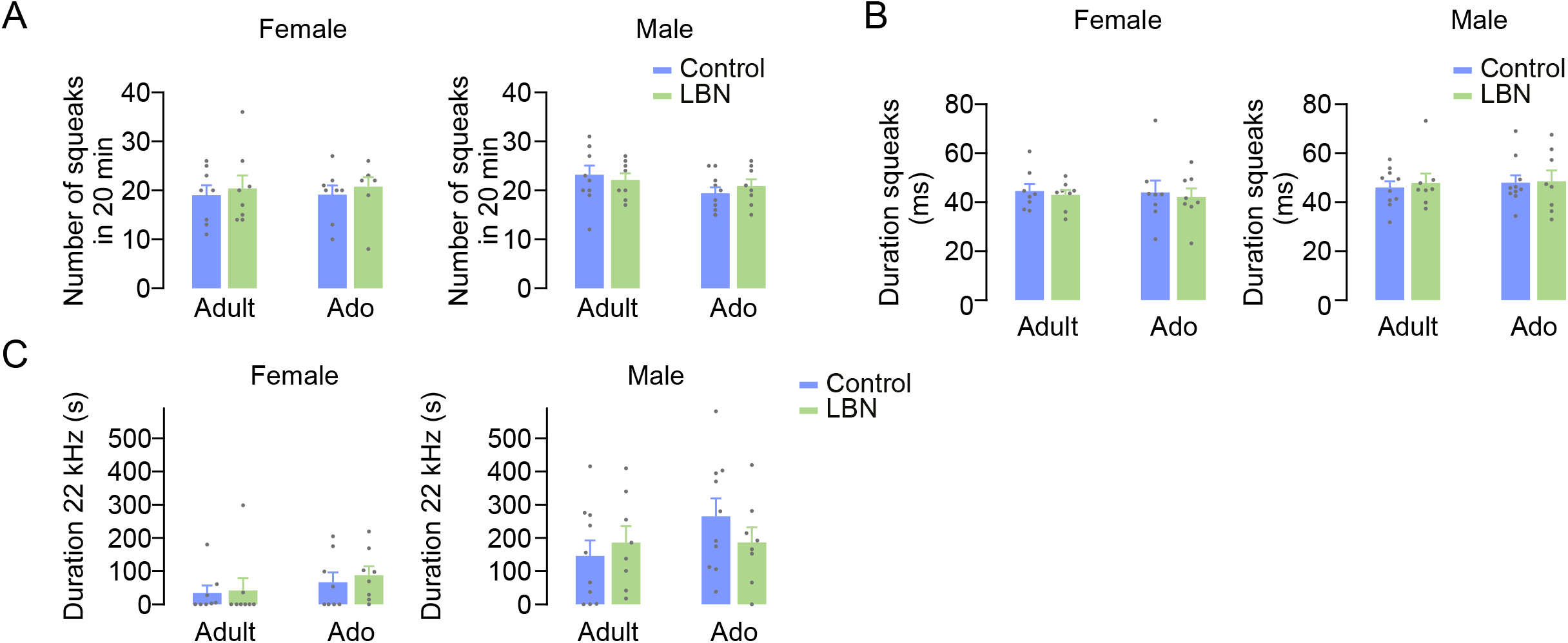
Squeaks and 22-kHz vocalizations during footshocks were not affected by either EAE or LAE. A. Number of squeaks in 20 minutes during the total shock period in control (adult n = 8, adolescent n = 8) and LBN (adult n = 8, adolescent n = 8) conditions for females (left) and males (right): control (adult n = 10, adolescent n = 10) and LBN (adult n = 8, adolescent n = 8) B. Average duration of squeaks during the total shock period in control and LBN conditions for females (left) and males (right). C. Average duration of 22-kHz vocalizations during the total shock period in control and LBN conditions for females (left) and males (right).

**Supplementary Method Figure 1. Timeline of handling and habituation for EC**

Day 1-5 Handling: Observers and demonstrators were handled for 5 minutes per day. Day 6 Habituation 1: First habituation session to the EC apparatus (10 min). Day 7 LAE manipulation. Day 8-9 Habituation 2 and 3: Second and third habituation sessions (10 min each). Day 10 EC test.

