## Supplementary figures and images for "Dissociable Effects of Early and Adolescent Adversity on Emotional Contagion"

### Supplemental Method figure 1

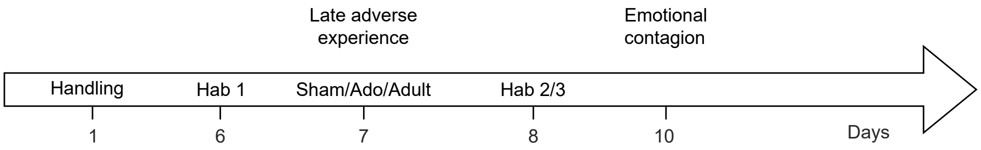
