## Supplemental material for "Dissociable Effects of Early and Adolescent Adversity on Emotional Contagion"

Supplementary Material.

**Subjects**

Primiparous Sprague Dawley dams (gestational day 15) were obtained from Janvier Labs and housed undisturbed in the animal facility until parturition. Dams were housed individually in standard cages within a quiet, temperature-controlled room under a 12:12 h reversed light/dark cycle, with food and water available ad libitum. They remained in these conditions throughout the remainder of pregnancy and until pup weaning. Demonstrator rats used in the EC test were also Sprague Dawley rats, acquired as adults from Janvier Labs. All demonstrators were naïve to footshocks and to the LBN condition, and were unfamiliar to the observer rats, though matched by sex and age. Demonstrators were delivered to the animal facility one week prior to testing to allow for acclimatization.

**Experimental procedures**

*LBN*

Dams were housed in standard Type IV rat cages (top outside dimensions: 590 x 380 mm; bottom inside dimensions: 530 x 330 mm; height: 200 mm; floor area 1815 cm^2^). Cages were provided with ¼-inch corn cob bedding (The Anderson) and paper towel (Scott Essential Multifold Paper Towels, white, 23.4 x 23.9 cm) as nesting material.

Births were monitored at least twice daily around the expected parturition date, with the day of birth designated as postnatal day (P) 0^30^. Litters remained undisturbed until P2 (morning), when pups from multiple litters were pooled and randomly assigned to the LBN or control conditions.

Litters were culled to 12 pups each, maintaining a 1:1 male-to-female ratio. Pups were briefly and gently removed from the cages, and sex was determined by assessing anogenital distance. Male and female pups were placed in separate, euthermic holding cages during the procedure. After weighing, pups were randomly assigned to LBN or control groups, with the order of assignment counterbalanced to minimize variability in separation duration. Total separation time was kept under 10 minutes. Litter mixing and redistribution were completed within a maximum of 12 hours across litters. To facilitate maternal acceptance and reduce stress from cage change, a small amount of the dam’s original bedding was transferred to the clean LBN or control cages and placed around the nest site.

In the control condition, cages were provided with approximately 550 gr of ¼-inch corn cob bedding and one trifold paper towel as nesting material. The bedding and nesting material were left unchanged throughout the procedure and remained in place until P9.

In the LBN condition, a custom-made metal mesh (grid size: 0.4 x 0.9 cm, triangle-shaped) was positioned 2.5 cm above the cage floor. Beneath the mesh, cages contained approximately 120 gr of ¼-inch corn cob bedding and half of a trifold paper towel as nesting material, limiting dam access to both nesting material and bedding.

Both control and LBN cages remained undisturbed from P2 to P9, with the exception of routine food and water replenishment. No bedding or nesting material was replaced during this period. On the morning of P9, all dams and pups were returned to clean, standard housing conditions with regular bedding and a full paper towel provided as nesting material.

*Weaning*

At weaning (P21), animals were housed in same-sex sibling pairs (one pair per cage). From this point onward, cages were enriched with basic supplements such as plastic tunnels to promote environmental stimulation.

*LAE procedure*

Each animal was placed individually in an inverted pyramid–shaped apparatus, wide at the top (450 × 450 mm) and narrow at the bottom (280 × 280 mm). A vanilla scent was introduced to differentiate this context from the later EC test. The session consisted of a 10-minute baseline period followed by a 20-minute shock period, during which four 1-second shocks (0.8 mA) were administered at random intervals of 4 or 6 minutes. Behavior was recorded using a video camera (Axis P1364 HDTV, Sweden), and an ultrasonic recording system (Avisoft UltraSoundGate 116H, Germany) coupled with a microphone (Avisoft CM16/CMPA-P48, Germany). The sampling rate was 250 kHz.

*Emotional contagion test*

EC tests were conducted in a two-chamber apparatus (length: 24 cm, width: 25 cm, height: 34 cm) (Med Associates, Fairfax, Vermont, United States), with a removable plexiglass partition containing vertical slits between chambers. The apparatus was housed within a light- and sound-attenuated cabinet and illuminated with infrared light. Observers were placed in a chamber with a perforated plastic floor, while demonstrators were placed in an adjacent chamber with a metal-grate floor for shock delivery. Each observer-demonstrator dyad was matched for age and sex. Tests included a 12-minute baseline phase, followed by a 12-minute shock phase during which the demonstrator received five 1-second footshocks (1.5 mA) at randomized intervals of 2 or 3 minutes. Behavior was recorded with a top-mounted video camera (Basler GigE acA1300–60gm, The Netherlands). Audio was captured using an ultrasonic recording system (UltraSoundGate 116H, Avisoft, Germany) coupled with one microphone (Avisoft CM16/CMPA-P48, Germany), placed at the top center of the chamber. The sampling rate was 250 kHz.

*Immunostainings and microscopy*

Rats were anaesthetized with an overdose of pentobarbital (60 mg/ml), 90 min after the EC test, and then transcardially perfused with phosphate-buffered saline solution (PBS) followed by ice-cold 4% paraformaldehyde (PFA) in PBS. Dissected brains were stored in 30% sucrose in PBS for cryoprotection before storage at -80°C. Coronal brain sections (40 μm thick) were cut using a CM3050 S cryostat (Leica Biosystems, USA), and every 6th section was collected in PBS. To assess neuronal activation in the area 24, immunofluorescent labeling was done for c-Fos, an immediate early gene, commonly used as marker of general neuronal activity. Briefly, to block the background and open the tissue matrix, free-floating sections were first incubated in 3% bovine serum albumin (BSA) and 0.3% Triton X-100 (Sigma, Cas. no.: 9036-19-5) in PBS at room temperature for 2 hrs, and then incubated with rabbit anti-c-Fos antibody (1:1000 dilution; Synaptic Systems: 226 003; RRID: AB_2231974) at 4 °C overnight. After washing three times with PBS, sections were incubated with Alexa Fluor 594 donkey anti-rabbit secondary antibody (1:800 dilution; Invitrogen: A21207; RRID: AB_141637) at room temperature for 2 hrs. Following three washes with PBS, sections were mounted with a mounting medium including DAPI (Vectashield Vibrance; Vector Laboratories: H-1800) onto glass microscopic slides and covered with glass cover slips. All images were acquired with a Zeiss Axio Scan Z1 fluorescent slide scanning microscope (software: Zeiss Zen 3.7; Carl Zeiss AG, Germany) at 10x magnification (Plan-Apochromat 10x/0.45 M27) and a resolution of 0.65 µm/pixel with a digital CMOS camera (ORCA-Flash4 V3; Hamamatsu Photonics K.K., Japan).

**Analysis**

*Maternal behaviors*

Scoring focused on two main behavioral categories: pup-directed behaviors, including licking and grooming, pup carrying, arched-back nursing, low nursing, and passive/side nursing, and non-pup-directed behaviors, such as self-grooming, time spent off-nest, rearing, and eating/drinking. The video-recordings were acquired at 10h00, during the dark phase of the light/dark cycle.

*Entropy*

The entropy index quantifies how unpredictable the transition from one behaviour to the next behaviour is for each dam. The entropy index for any one dam ranges from 0, where behaviour is completely predictable, with any given behavior i always followed by a particular behavior j (i.e. pij=1), whilst all other transitions have probability zero, to the logarithm in base 2 of the number of behaviours analysed here (nest building, tail chasing, licking/grooming, arched-back nursing, side/passive nursing, low nursing, carrying pups, selfgrooming, off-nest, move around nest, eat/drink, rearing), log2(12)=3.58, where all transitions are equiprobable and the dam’s behavioural transitions are maximally entropic, or unpredictable. Furthermore, an additional level of complexity is added to the entropy calculation by using the empirical transition matrix; as described by Demaestri et al. (2022) and Molet et al. (2016): entropy is calculated for each row, each of which represents a probability distribution, and then combined into a singular measure. The stationary distribution πi of the Markov chain (a measure of long-term probability of each behaviour) is calculated and used to weight the entropy values. The entropy index is calculated with the equation below, where *i* represents the initial behaviour, *j* represents the following behaviour, and P*ij* represents the probability of the transition from the initial behaviour (*i*) to the following behaviour (*j*). Given that LBN was imposed from P2 to P9, to assess whether LBN altered entropy, we directly compared entropy from P3 to P7 (sampled at P3, P5 and P6-7).

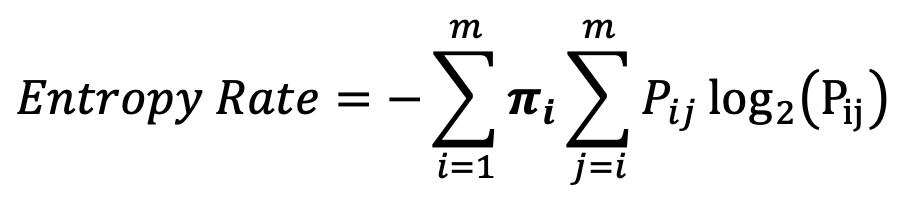

*Vocalizations*

The “Long Rat call v2” network was applied for detection of 22-kHz vocalizations using a frequency range of 18–100 kHz. 50-kHz vocalizations were detected using a high-frequency cutoff of 100 kHz and a low-frequency cutoff of 30 kHz, using the “All short calls v1” network. False positives were manually rejected. Squeaks were identified through visual inspection of the spectrogram. The frequency of 50-kHz vocalizations was extracted from the spectral band corresponding to the peak intensity of each vocalization and quantified as the mode of its distribution, given its skewed nature (Supplementary figure 1B and C).

*Pose estimation*

The following body parts were labeled: “Nose", "Right_before_eye", "Right_before_ear", "Right_after_ear", "Left_before_eye", "Left_before_ear", "Left_after_ear", "Skeleton_1", "Skeleton_2", "Skeleton_3", "Tail", "Right_side_3", "Right_side_tail", "Left_side_3", and "Left_side_tail”. Labels with a DeepLabCut likelihood score below 0.6 were discarded. Box coordinates (i.e., platform corners) were manually extracted in Fiji from each video. These coordinates, along with body part labels, were then converted and merged into a common spatial coordinate plane to reconstruct the original setup as if videos had not been cropped. For each frame, we computed average positions for the head, body, and tail of each animal. Metrics were then calculated frame-by-frame, including (1) the distance of the observer’s head to the separator between the observer and the demonstrator, and (2) the angle between the observer’s and demonstrator’s head directions. To minimize overestimation caused by rearing behaviors (where body parts extend beyond the platform), we projected all body parts within the horizontal plane of the platform before computing distances. Finally, all distance-based measures were normalized to the diagonal length of the platform box, constraining values to approximately the [0,1] interval.

*Cell counting*

At first, random sections from random animals were selected to find the optimal quantification parameters (e.g., threshold, cell size), then all sections were analyzed using the same parameters. Results were checked by a researcher to avoid misquantification. Samples with poor tissue or staining quality, extensive damage, or imaging issues were excluded from the analysis. c-Fos counts were summed up per region per animal and normalized to the total analyzed area (mm^2^).

**Statistical analysis**

The number of animals, the Bayes factor, and the test used for each analysis are specified in the Results section. For datasets with *n* ≥ 6 normality was assessed using the Shapiro-Wilk test. If the data were normally distributed, independent samples *t*-tests were used for two-group comparisons. For non-normally distributed data or small sample sizes (*n* < 6), the non-parametric Mann–Whitney U test was used for group comparisons. Data involving more than two groups were analyzed using two- or three-way ANOVAs, following an inspection of Q–Q plots to assess normality and homogeneity of variances. The mediation analysis was implemented in JASP (method: robust) in order to estimate whether vocalization characteristics (ratio between the average rate during baseline and shock periods of 50 kHz and 22k-Hz rates) or observer immobility (expressed in percentage) could mediate the effect of LBN on the Demonstrator immobility during shock experience.
